# CSRefiner: A lightweight framework for fine-tuning cell segmentation models with small datasets

**DOI:** 10.1101/2025.09.14.675731

**Authors:** Can Shi, Yumei Li, Jing Guo, Qiuling Chen, Tingting Cao, Sha Liao, Ao Chen, Mei Li, Ying Zhang

**Author notes:** Correspondence address. Mei Li and Ying Zhang, BGI Research, Shenzhen 518083, China.

## Abstract

Recent advances in spatial omics technologies have enabled transcriptome profiling at subcellular resolution. By performing cell segmentation on nuclear or membrane staining images, researchers can acquire single cell level spatial gene expression data, which in turn enables subsequent biological interpretation. Although deep learning-based segmentation models achieve high overall accuracy, their performance remains suboptimal for whole-tissue analysis, particularly in ensuring consistent segmentation accuracy across diverse cell populations. Existing fine-tuning approaches often require extensive retraining or are tailored to specific model architectures, limiting their adaptability and scalability in practical settings. To address these challenges, we present CSRefiner, a lightweight and efficient fine-tuning framework for precise whole-tissue single-cell spatial expression analysis. Our approach incorporates support for fine-tuning widely uaed segmentation models in the field of spatial omics, including recent published model Cellpose-SAM, while achieving high accuracy with very limited annotated data. This study demonstrates CSRefiner’s superior performance across various staining types and its compatibility with multiple mainstream models. Combining operational simplicity with robust accuracy, our framework offers a practical solution for real-world spatial transcriptomics applications.

## 1 Introduction

Spatially resolved transcriptomics (SRT) technologies have opened a new door for investigating the cellular microenvironment and cellular heterogeneity in complex tissues[1, 2, 3, 4, 5], these technologies were named the “Method of the Year 2020” by *Nature Methods*[6]. Recent advances in spatial omics now enable RNA capture at subcellular resolution, as demonstrated by platforms such as Stereo-seq[7] (0.5 *µ*m), Visium HD[8] (2 *µ*m), and Seq-Scope[9] (0.5–0.8 *µ*m). These high-resolution SRT methods enable researchers to investigate single-cell transcriptomes, unlocking insights into functional, developmental, and disease-related mechanisms of living organisms[1]. To obtain single-cell data, cell segmentation serves as the core step for extracting single nuclear or cell boundaries from cell staining images[10, 11, 12, 13, 14]. Those single-cell analysis pipelines in spatial transcriptomics (SRT) typically employ state-of-the-art deep learning models for cell segmentation, such as Cellpose[15, 16, 17, 18], StarDist[19], and the U-Net-based cell segmentation models[20].

Although current cell segmentation algorithms exhibit satisfactory performance under controlled conditions, their practical application encounters significant challenges due to cellular morphological heterogeneity and variations in image quality[14, 20, 21, 22]. Specifically in entire sections of complex biological tissues that contain multiple cell types, these algorithms often fail to maintain consistent segmentation accuracy across all cellular populations (see Results and Fig. 1A). Even if the model achieves a segmentation accuracy of more than 95% in the global evaluation, under-segmentation or over-segmentation of local areas may still lead to key segmentation errors, which will further affect the reliability of differential expression analysis, identification of rare cell subpopulations, and construction of cell spatial interaction networks, and even lead to erroneous biological conclusions. Manual annotation is often the ultimate solution for cell segmentation, but large or multiple samples will bring more manpower. Annotating a single 1 cm × 1 cm sample often takes an annotator half a month or even longer. This high-intensity manual effort not only greatly increases the research cost, but also seriously restricts the research progress.

**Figure 1:**
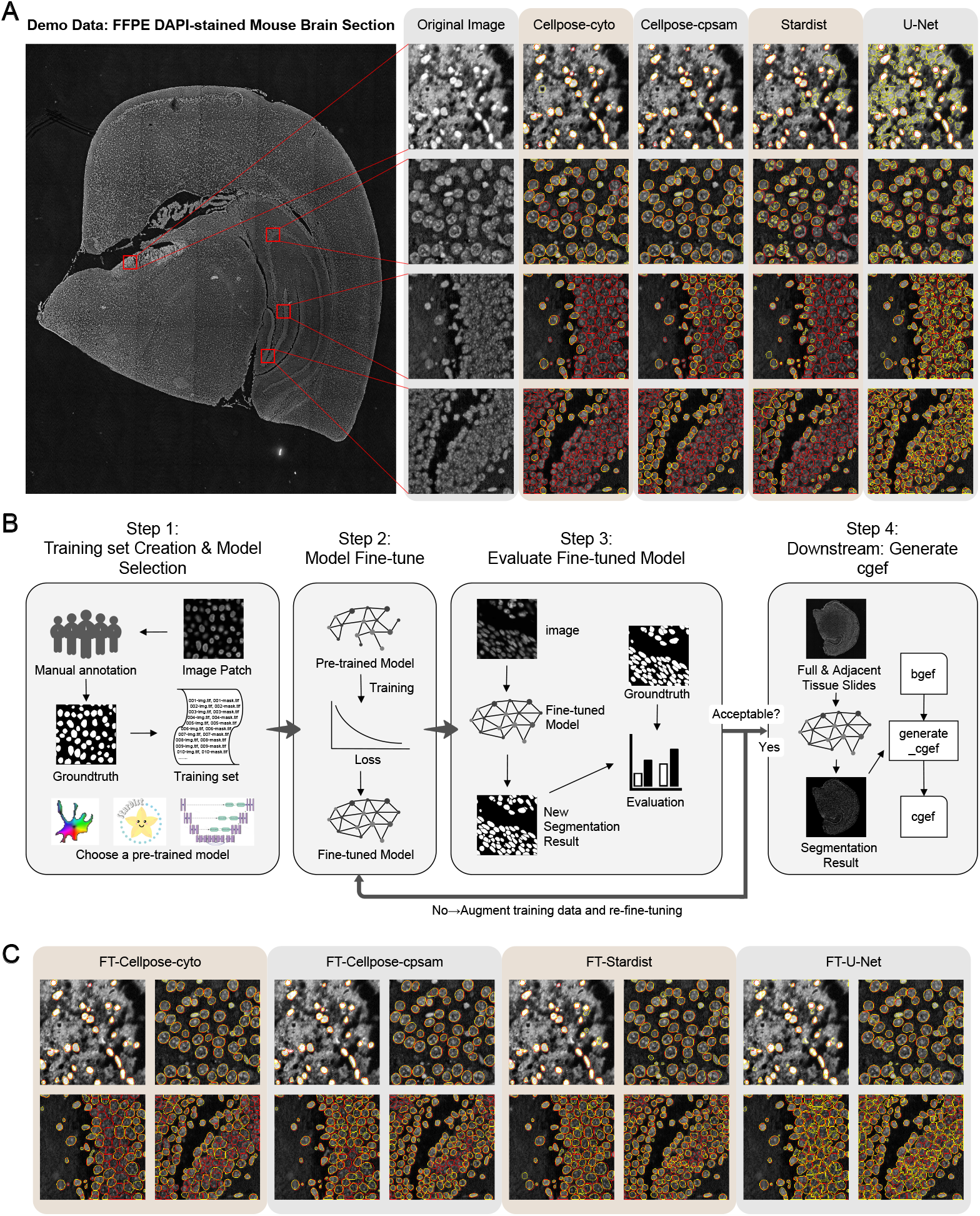
Performance of pre-trained and fine-tuned segmentation models on FFPE DAPI-stained mouse brain section, and schematic workflow of CSRefiner. **Description for Figure 1:** (A) Representative segmentation results from four pre-trained models (Cellpose-cyto, Cellpose-cpsam, StarDist-2D_versatile_fluo, and U-Net) for FFPE mouse brain tissue with DAPI-stained. Four regions are shown, including hippocampus and non-hippocampal areas. Red contours indicate ground-truth manual annotations, and yellow contours indicate model-predicted segmentation boundaries. (B) Schematic overview of the CSRefiner workflow, including training set preparation, model selection and fine-tuning, performance evaluation, and downstream generation of cgef files. (C) Segmentation results after fine-tuning with CSRefiner, corresponding to the regions shown in panel A. Model names prefixed with “FT-” indicate fine-tuned versions (same hereafter).

In the face of the above challenges, we developed CSRefiner, a lightweight and efficient fine-tuning framework compatible with multiple cell segmentation models, designed to obtain single-cell spatial expression data with accurate whole-tissue segmentation results. By introducing a small sample fine-tuning mechanism, the framework can quickly adapt the existing model to specific image features, thereby significantly improving the segmentation accuracy. Currently, it is compatible with three mainstream cell segmentation models in the field of spatial transcriptomics, namely Cellpose, StarDist, and a U-Net-based cell segmentation model in CellBin—a generalist framework for constructing single-cell expression matrices in spatial omics (hereafter referred to as U-Net), and will be further expandable to support more models in the future. Additionally, it is applicable to a variety of staining methods, such as hematoxylin and eosin (H&E), single-stranded DNA (ssDNA), 4’,6-diamidino-2-phenylindole (DAPI), and multiplex immunofluorescence (mIF). To facilitate evaluation and benchmarking, we also generated and publicly release two demo datasets consisting of formalin-fixed, paraffin-embedded (FFPE) DAPI-and fresh frozen (FF) H&E-stained mouse brain sections with paired expression matrices and annotation results. Collectively, these capabilities enable CSRefiner to significantly minimize manual intervention while preserving segmentation accuracy, particularly in critical regions. This synergistic enhancement not only accelerates the spatial transcriptomics analysis pipeline but also unlocks the potential for robust, large-scale investigations by providing researchers with a scalable and cost-effective annotation strategy.

## 2 Results

### 2.1 CSRefiner framework overview

Before developing CSRefiner, we first evaluated the four segmentation models supported by CSRefiner for fine-tuning on FFPE DAPI-stained and FF H&E-stained mouse brain sections. Visual inspection of segmentation results (Fig. 1A, Supplementary Fig.1) revealed obvious performance inconsistencies across tissue regions: in non-hippocampal areas with sparse cells and clear nuclear boundaries, models generally captured cell outlines; however, in the hippocampus (dense cells, blurred nuclear boundaries), all pre-trained models had notable defects.

This inability of mainstream models for SRT to maintain visually reliable segmentation in complex tissue regions (e.g., the hippocampus) motivated us to develop CSRefiner. As illustrated in Fig. 1B, CSRefiner is a lightweight and flexible pipeline designed to rapidly adapt pre-trained segmentation models to tissue-specific features in spatial transcriptomics images. The workflow consists of four streamlined steps. First, users construct a small training set by annotating representative patches from regions where baseline segmentation is suboptimal, such as densely packed or morphologically heterogeneous regions (e.g., hippocampus). A base model is then selected from supported models (Cellpose, StarDist, or U-Net). Second, the chosen model is fine-tuned on the annotated patches through a unified, parameter-controlled script. Third, the refined model is evaluated on the training set/testing set and iteratively refined by adding more annotated patches if necessary, ensuring that the model achieves satisfactory performance across challenging regions. Finally, the optimized model is applied to whole-slide images, and the resulting segmentation boundaries are integrated with the corresponding expression matrix to generate a single-cell-level gene expression format (cgef) file, a widely used standard in spatial transcriptomics, facilitating downstream single-cell analyses.

### 2.2 CSRefiner improves segmentation accuracy with minimal manual effort

#### 2.2.1 Improvement of segmentation performance

In non-hippocampal regions, CSRefiner achieves more refined segmentation, closely adhering to actual cell nucleus outlines and rarely missing cells (Fig. 1C). U-Net, which previously frequently produced false positives in the background or over-segmented cells, now generates complete cell outlines, significantly reducing the fragmentation of individual nuclei into disjoint fragments. In the hippocampus, where dense cell packing and blurred nucleus boundaries lead to severe false negatives in pre-trained models, CSRefiner achieves significant improvements: it successfully recovers many previously overlooked cells and can distinguish individual nuclei even in the densest clusters. Although U-Net’s boundary delineation accuracy remains slightly lower than Cellpose and StarDist (consistent with its inherent emphasis on recall), its segmentation results are visually much more consistent than the baseline models.

We next assessed the impact of CSRefiner on segmentation accuracy using five standard evaluation metrics: precision, recall, F1 score, Jaccard index, and Dice coefficient (Fig. 2A–E). Boxplot comparisons showed improvements across all metrics for all tested models. Cellpose-cyto F1 increased from 0.54 to 0.78, Cellpose-cpsam from 0.79 to 0.91, StarDist from 0.42 to 0.87, and U-Net from 0.37 to 0.71. Similar gains were observed for precision, recall, Dice, and Jaccard indices. Notably, the improvement was particularly pronounced in models with weaker initial performance, such as StarDist and U-Net, while even the high-performing Cellpose-cpsam model achieved measurable gains. Moreover, the reduced variance in post-fine-tuning scores indicates enhanced consistency and robustness of the models across different image regions.

**Figure 2:**
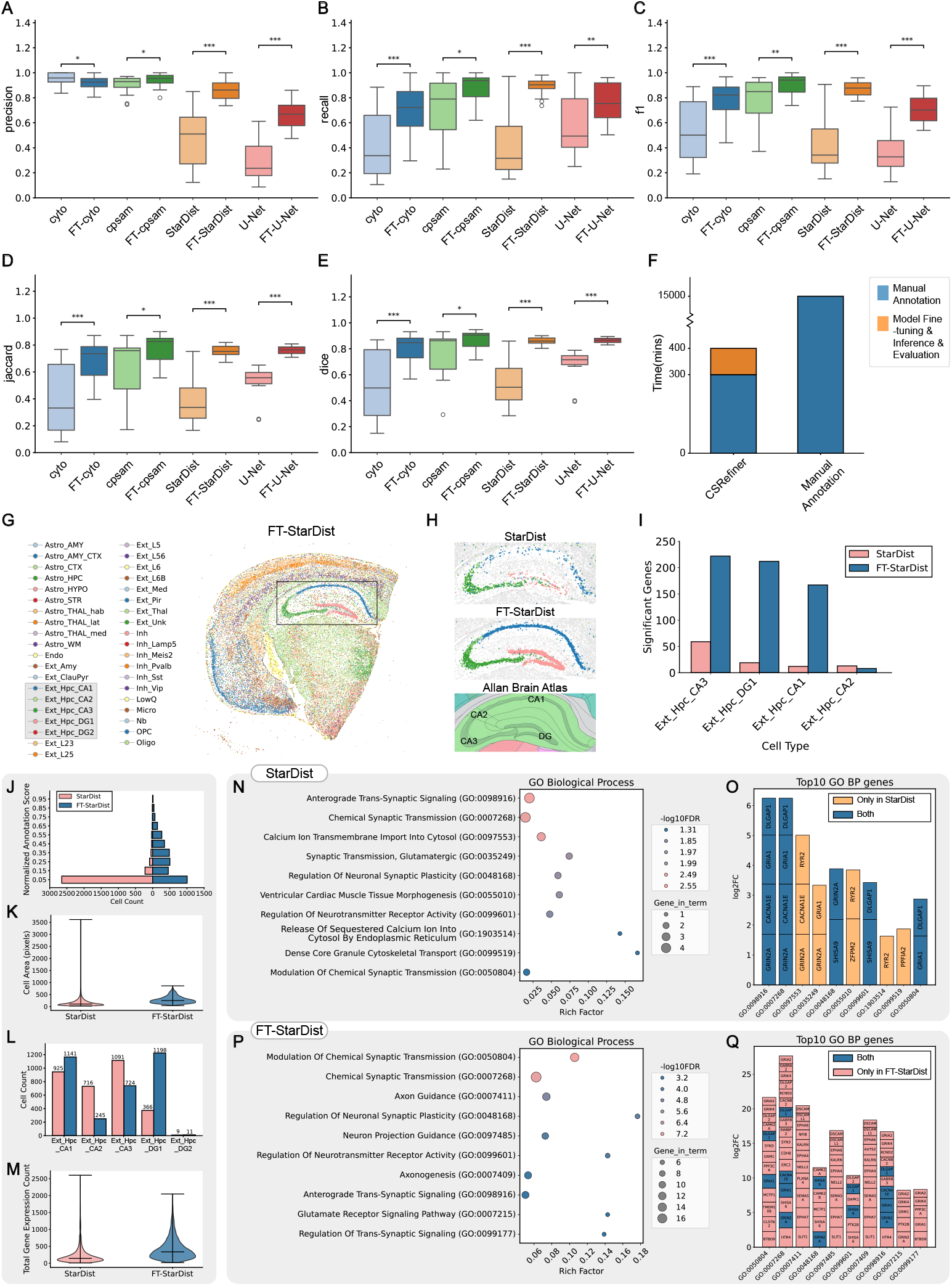
Segmentation performance and downstream biological analyses improved by CSRefiner. **Description for Figure 2:** (A–E) Quantitative evaluation of segmentation performance before and after fine-tuning across four representative models (Cellpose-cyto, Cellpose-cpsam, StarDist, and U-Net). Boxplots show improvements in (A) precision, (B) recall, (C) F1 score, (D) Jaccard index, and (E) Dice coefficient. The significance mark line and P value are added to the box plot. The number of “*” from 1 to 3 represents a P value less than 0.05, 0.01, and 0.001. (F) Time required for manual whole-slide annotation ( 10 days) versus CSRefiner-assisted workflow ( 400 minutes). (G) Spatial maps of cell type annotations generated by cell2location using cgef matrices from the fine-tuned StarDist model. (H) Visualization of segmented and annotated cells in hippocampal subregions before and after fine-tuning compared with the Allan Brain Atlas for the hippocampus. (I) Comparison of the number of significant genes in each cell subtype in the hippocampus before and after fine-tuning. (J) Distribution of normalized annotation scores in the hippocampus before and after fine-tuning. (K) Distribution of cell areas in the hippocampus before and after fine-tuning. (L) Comparison of the number of cells of each cell subtype in the hippocampus before and after fine-tuning. (M) The number of genes detected per cell in the hippocampus before and after fine-tuning. (N-Q) Gene ontology enrichment analysis results for Ext_Hpc_DG1 before and after fine-tuning. In the bubble plot, bubbles positioned further right indicate stronger enrichment, with redder color showing higher significance and larger size representing more hit genes. The stacked bar plot displays contributing genes for each GO term, where bar height reflects contribution strength (taller bars highlight potential driver genes). Orange denotes terms/genes present only before fine-tuning, blue indicates both, and pink marks those unique to after fine-tuning.

#### 2.2.2 CSRefiner saves more time than manual annotation

CSRefiner greatly reduced annotation time. We compared the overall time required for CSRefiner-assisted fine-tuning against full manual annotation (Fig. 2F). Based on an average annotation time of 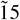 minutes per 256 × 256 image patch, generating 20 training patches required 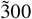 minutes. Model fine-tuning, evaluation, and inference added an additional 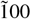 minutes, resulting in a total of 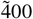 minutes (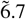 hours) for a complete CSRefiner workflow. In contrast, manual annotation of an entire whole-slide image would require approximately 10 days (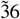 × longer) under comparable conditions. These results demonstrate that CSRefiner enables the generation of high-quality segmentation results within a fraction of the time, significantly lowering the cost and labor of manual annotation.

### 2.3 Downstream biological impact of improved segmentation

To further evaluate the biological relevance of CSRefiner, we examined how improved segmentation accuracy influenced downstream spatial transcriptomics analyses.

#### 2.3.1 Improved spatial mapping

Using StarDist, the model with the greatest performance gains, fine-tuned segmentation enabled the generation of high-quality cgef matrices. Subsequent cell type annotation with cell2location[23] produced spatial distributions that closely matched established anatomical structures and biological priors (Fig. 2G). Notably, hippocampal subregions were delineated with high fidelity (Fig. 2H), confirming that CSRefiner-generated segmentations directly support biologically meaningful spatial mapping.

#### 2.3.2 Improved DEGs detection and annotation quality in the hippocampus

Differential expression analysis of hippocampal cell types revealed that fine-tuning increased the number of significant genes in Ext_Hpc_CA3/DG1/CA1, consistent with visual inspection, indicating improved segmentation of dense hippocampal regions (Fig. 2I).

We then analyzed in detail the changes in cell annotation in the hippocampus before and after fine-tuning. First, annotation confidence scores shifted toward higher values, indicating more reliable cell type assignments (Fig. 2J). Quantitative assessment of cell morphology revealed that extreme cell size outliers were eliminated, while total cell area slightly increased, reflecting more regular and biologically plausible segmentation boundaries (Fig. 2K). Furthermore, cell type-specific counts exhibited biologically consistent changes (Fig. 2L): Ext_Hpc_CA1 and Ext_Hpc_DG1 cells increased significantly, Ext_Hpc_DG2 remained stable, and Ext_Hpc_CA2/CA3 decreased slightly. These results align with expected patterns of hippocampal cytoarchitecture. Finally, fine-tuning raised the number of detected genes per cell (Fig. 2M), thereby enhancing the sensitivity of downstream transcriptomic analyses. By recovering expression signals from cells that would otherwise be under-segmented or omitted, CSRefiner improved both the resolution and the biological interpretability of spatial transcriptomics data.

#### 2.3.3 Enables deeper biological discoveries

As shown in Fig. 2N–Q, we performed enrichment analysis of Ext_Hpc_DG1 using the GO Biological Process 2023 database[24, 25]. The enrichment analysis quality of the fine-tuned model significantly outperformed that of the pre-trained model. This difference stems from a more accurate cell segmentation mask that ensures the purity of gene expression signals in the cell bin matrix, ultimately outputting enrichment results that are more consistent with the true biological state.

There are significant overlaps in neural/synaptic pathways between the pre- and post-fine-tuning results, such as Chemical Synaptic Transmission (GO:0007268), Modulation of Chemical Synaptic Transmission (GO:0050804), Regulation of Neuronal Synaptic Plasticity (GO:0048168), Regulation of Neurotransmitter Receptor Activity (GO:0099601), and Anterograde Trans-Synaptic Signaling (GO:0098916). Before fine-tuning, this analysis highlighted a limited set of synaptic signaling processes, such as anterograde trans-synaptic signaling (GO:0098916) and chemical synaptic transmission (GO:0007268), supported by a small group of canonical synaptic genes (e.g., GRIA1, GRIN2A, CACNA1E, DLGAP1). Furthermore, the pre-trained top 10 included terms such as Ventricular Cardiac Muscle Tissue Morphogenesis (GO:0055010), which are inconsistent with brain/hippocampal biology, suggesting possible false positive enrichment due to the inclusion of non-target cells/background or noise in the differentially expressed gene list. By contrast, after fine-tuning, the same analysis revealed a broader and biologically coherent network of pathways, including axon guidance, neuron projection guidance, and axonogenesis (Fig. 2P). The expanded gene set (e.g., EPHA4/6/7, SLIT1, DSCAM/DSCAML1, KALRN, CAMK2A/B, GRIA1/2, GRIN2A) is shown in Fig. 2Q and represents critical regulators of axonal connectivity and synaptic plasticity, in line with the known functions of hippocampal excitatory neurons[26, 27, 28, 29].

### 2.4 Variable training sample needs for fine-tuning

To evaluate the required training set size for regions of varying segmentation difficulty, we systematically varied the number of annotated cells used for fine-tuning and tested on independent held-out regions not included in the training set.

Fig. 3A shows that in the hippocampus, model performance improved gradually with increased training examples: F1 scores rose from 0.23 with no training cells, to 0.35 with 105 cells, 0.36 with 357 cells, and 0.49 with 1,430 cells. Accuracy and recall showed similar upward trends, and the performance curve did not plateau at 1,430 cells, suggesting further gains with additional annotations. In contrast, in non-hippocampal regions, performance improved rapidly with relatively few training examples: F1 increased from 0.33 with no training cells, to 0.67 with 105 cells, 0.84 with 357 cells, and 0.85 with 566 cells. Here, the performance curve approached saturation after 300 cells, and additional annotations provided diminishing returns (Fig. 3B). These observations suggest that regional complexity critically determines training requirements: dense and heterogeneous areas such as the hippocampus require larger training sets to approach optimal performance, whereas simpler regions converge with fewer annotations.

**Figure 3:**
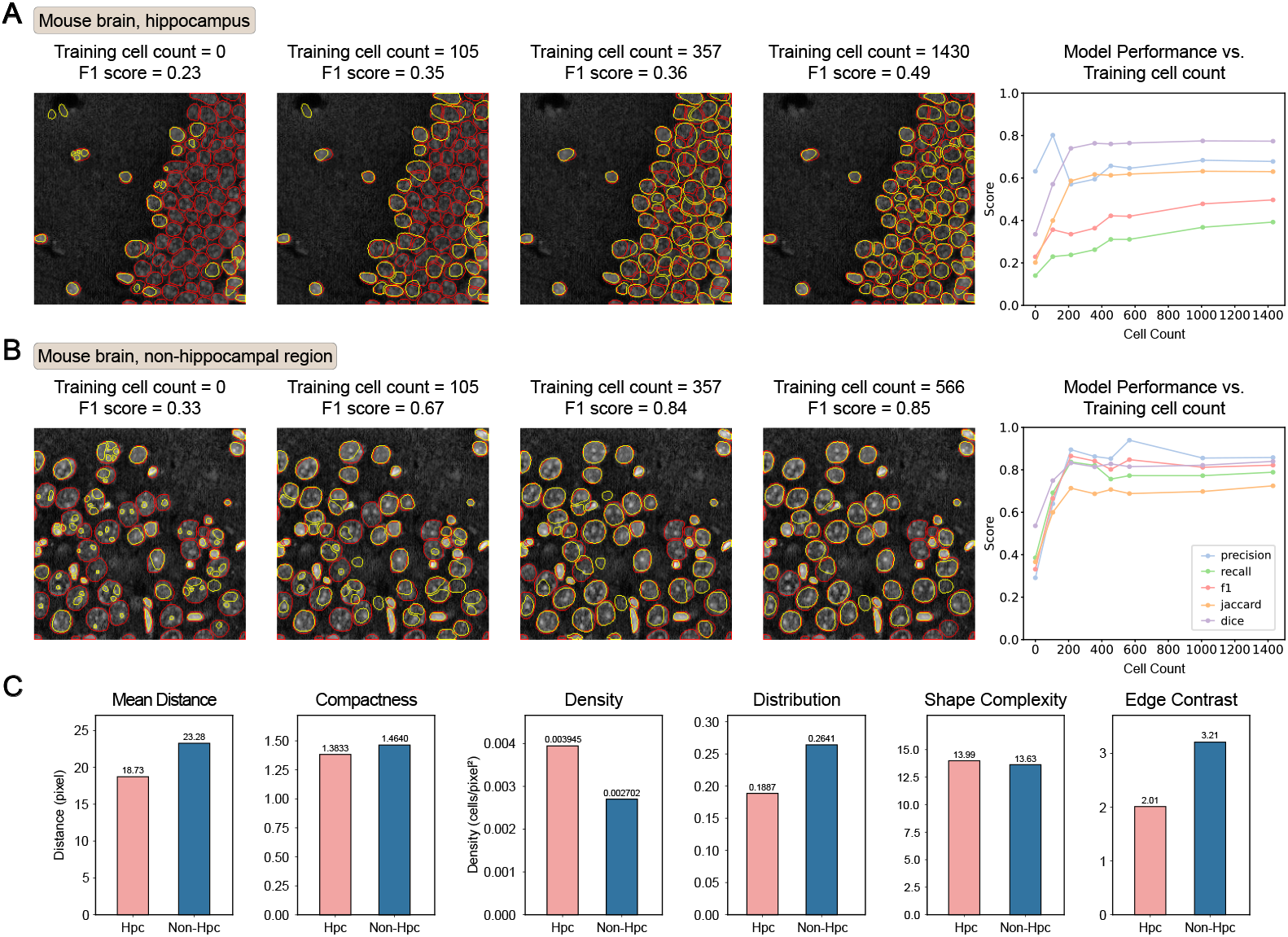
Training set size and tissue complexity influence segmentation performance. **Description for Figure 3:** (A) Model performance on the hippocampus (not included in training) after fine-tuning the model using different numbers of annotated cells. The line graph on the right visualizes precision, recall, and F1 score for different numbers of training cells. (B) Same as panel A, model performance on non-hippocampal regions (not included in training). (C) Quantitative comparison of regional structural complexity between hippocampal and non-hippocampal images. Metrics include mean distance between cells, compactness, density, distribution, shape complexity, and edge contrast.

To quantitatively assess region-specific complexity, we compared six structural metrics between hippocampal and non-hippocampal images (Fig. 3C). Hippocampal cells exhibited smaller mean inter-cell distance (18.7 vs. 23.3) and lower compactness (1.38 vs. 1.46), indicating tighter packing. Cell density was higher (0.0039 vs. 0.0027), while distribution uniformity was lower (0.189 vs. 0.264), suggesting large-scale dense clustering rather than localized groups. Edge contrast was reduced (2.01 vs. 3.21), reflecting weaker boundaries that complicate segmentation. In contrast, shape complexity was comparable between regions (13.99 vs. 13.63), implying that cell morphology was not a major differentiating factor.

Overall, these results confirm that hippocampal tissue poses greater segmentation challenges due to denser cellular organization and lower boundary clarity, consistent with qualitative visual inspection. Optimal training set design should therefore consider both region type and structural complexity, balancing annotation effort with expected performance gains.

## 3 Discussion

Accurate cell segmentation is fundamental to spatially resolved transcriptomic analysis, as errors at this stage propagate to downstream tasks such as cell type annotation, differential expression analysis, and pathway enrichment. In this study, we present CSRefiner, a flexible framework designed to fine-tune compatible cell segmentation models. Comparative evaluation demonstrates that CSRefiner substantially improves segmentation accuracy across multiple architectures, reduces annotation time by more than an order of magnitude, and enhances biological interpretability of downstream analyses. We also provide two high-quality datasets containing paired images, annotations, and expression matrices as community resources for benchmarking and method development.

A key finding is that model fine-tuning tailored to tissue-specific features markedly outperforms direct application of pre-trained models. In challenging regions such as the hippocampus, fine-tuned models achieved notable gains in segmentation metrics, recovering cells that were otherwise missed. These improvements translated into more coherent spatial maps and more biologically consistent pathway enrichment results. Beyond tissue structural complexity, another major challenge lies in the intrinsic properties of FFPE samples. Formalin fixation and long-term storage cause nucleic acid fragmentation and degradation, while nonspecific dye binding elevates background noise and reduces boundary clarity, making segmentation particularly difficult. Remarkably, CSRefiner achieved consistent gains on our FFPE demo datasets, even though the U-Net baseline had been trained on FF samples, highlighting the framework’s robustness across preparation protocols.

Our results further highlight the importance of training set design. Performance scaling analyses revealed that simpler regions reach saturation with relatively few annotated cells, whereas complex structures such as the hippocampus require substantially larger training sets to achieve comparable accuracy. Quantitative metrics of tissue organization—such as cell density, compactness, and boundary clarity—correlated well with segmentation difficulty, providing a principled way to estimate annotation requirements for different tissue contexts. These insights may inform annotation strategies in future large-scale studies, enabling an optimized balance between manual effort and segmentation performance.

While CSRefiner reduces annotation requirements compared with full retraining, some manual effort remains necessary, especially in highly complex tissues. In addition, integration with multimodal spatial data, such as joint proteomic profiling, may require further adaptation. Future extensions could incorporate interactive annotation tools to streamline training set preparation and broaden compatibility with emerging segmentation backbones.

## 4 Methods

### 4.1 Tissue samples and data acquisition

#### 4.1.1 Experimental animals and ethical approval

Two mouse brain tissue sections were used in this study. Brain tissues were collected from 6-week-old C57BL/6 male mice, provided by the Guangzhou Institutes of Biomedicine and Health, Chinese Academy of Sciences, in collaboration with the BGI laboratory. All animal experiments followed institutional ethical guidelines and were approved by the relevant committees.

#### 4.1.2 Tissue preparation and staining

FFPE DAPI-stained mouse brain section: Fresh mouse brains were dissected and fixed in 4% paraformaldehyde (Servicebio, G1121) at room temperature for 24-48 hours, followed by dehydration, paraffin embedding, and sectioning into 5 *µ*m-thick slices using standard histopathological protocols. Sections were mounted on Stereo-seq Chip N after flattening in a 45°C water bath, then baked at 42°C for 3 hours and dried overnight. Deparaffinization, decrosslinking (95°C with FFPE Decrosslinking Reagent; STOmics, 211KN114-EA), and post-decrosslinking fixation (−20°C methanol) were performed according to the STOmics Stereo-seq Transcriptomics Set protocol[30]. DAPI staining (1 *µ*m/mL, room temperature, 10 minutes in the dark) was conducted after washing with 0.1×SSC buffer (Thermo Fisher Scientific, AM9770) supplemented with RNase inhibitor (GCATBio, LS-EZ-E-00006Q).

FF H&E-stained mouse brain section: Dissected mouse brains were snap-frozen in dry ice-cooled Tissue-Tek OCT (Sakura, 4583) and sectioned into 10 *µ*m-thick coronal slices using a Leica CM1950 cryostat. Sections were mounted on Stereo-seq Chip N, incubated on a 37°C slide dryer for 5 minutes, and fixed in −20°C methanol for 30 minutes. Adjacent sections on glass slides were stained with H&E using a Solabio G1121 kit (hematoxylin for 8 minutes, eosin for 3 minutes) following the manufacturer’s instructions, then dehydrated, cleared, and mounted with neutral resin.

#### 4.1.3 Imaging data acquisition

FFPE DAPI-stained mouse brain section: Fluorescence images were acquired using a STOmics Microscope Go Optical system (Scanner Version 1.2.2, 20×/0.5 NA objective, Ximea Mc124 camera) in epi-fluorescence mode (excitation: 358 nm, emission: 461 nm). Image quality was verified via the StereoMap software (imageQC module).

FF H&E-stained mouse brain section: Brightfield images were captured using a Motic PA53 FS6 microscope (PA53Scanner 1.0.0.14, 10×/0.75 NA objective, PA53 FS6 SCAN S5LITE RGB module) in epi-brightfield mode, with images saved in TIFF format.

#### 4.1.4 Gene expression matrix generation

FFPE and FF sections on Stereo-seq chips were processed for RNA capture, reverse transcription, and cDNA amplification following the STOmics Stereo-seq Transcriptomics Set protocol[30]: FFPE samples used random primers for total RNA capture and FFPE Decrosslinking Reagent for decrosslinking, while FF samples underwent methanol fixation and short-time permeabilization. Libraries were constructed via cDNA fragmentation (Tn5 transposase) and PCR, then sequenced on MGI DNBSEQ-T1 (FFPE) or DNBSEQ-Tx (FF) sequencers.

Raw sequencing data were processed with the SAW pipeline[31]: low-quality reads were filtered, CID sequences were mapped to chip coordinates (1 base mismatch allowed), and reads were aligned to the mm10 genome (STAR software[32], MAPQ > 10). UMIs for the same CID and gene were collapsed to eliminate PCR duplicates, and spatial gene expression matrices were generated by integrating gene counts with coordinates.

### 4.2 Model selection, patch generation, and training set construction

To enable efficient fine-tuning with limited annotated data, CSRefiner encourages users to construct a compact yet informative training set from the target images. The process begins with the selection of a suitable pre-trained model. Specifically, users are advised to apply the available pre-trained segmentation models supported by CSRefiner (e.g., Cellpose, StarDist, or U-Net) to the target tissue image and visually inspect the initial segmentation. This assessment, typically carried out using widely adopted visualization tools such as ImageJ, involves evaluating both global coverage and local boundary accuracy. The model that demonstrates the most satisfactory overall performance is then chosen as the starting point for fine-tuning.

Following model selection, the full tissue image is divided into smaller patches for training. CSRefiner provides a lightweight cropping utility that supports batch processing of whole-slide images and allows users to customize patch sizes to facilitate this step. With the exception of StarDist’s H&E-optimized pre-trained model (‘2D_versatile_he’), which requires training patches of 512 × 512 pixels, all other models operate with patches of 256 × 256 pixels. For convenience, CSRefiner also includes a padding utility that can extend 256 × 256 images to the 512 × 512 size without altering cell morphology. This is achieved by symmetrically reflecting the image content at the borders (border reflection padding), which preserves the original cellular structures while avoiding artificial distortions introduced by resizing or interpolation. This ensures compatibility with different base models while avoiding the need to separately generate patches of multiple resolutions.

When selecting patches for annotation, priority should be given to regions where segmentation quality is noticeably suboptimal—for example, areas with blurred cell boundaries, overlapping or densely packed cells, or uneven staining that leads to under-segmentation or over-segmentation. To maintain model generalizability, a small proportion of morphologically simple and sparsely populated regions should also be included, although overrepresentation of these regions is discouraged to prevent model forgetting effects.

Annotation of the selected patches can be performed using any standard software familiar to the user. In practice, QuPath is recommended for its accessibility and compatibility. CSRefiner provides a step-by-step tutorial for generating cell annotations with QuPath. However, QuPath typically produces semantic masks (all cells assigned a common label), whereas instance segmentation models such as Cellpose and StarDist require instance masks, in which each cell is assigned a unique integer identifier. To address this mismatch, CSRefiner offers a conversion script that automatically transforms semantic masks into instance masks, ensuring training compatibility.

Finally, CSRefiner organizes the annotated images and corresponding masks into a standardized training file list. This file management system simplifies subsequent fine-tuning, enables reproducible execution of the workflow, and facilitates version tracking of training datasets. Based on empirical results, we recommend preparing at least 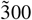 annotated cells to achieve satisfactory fine-tuning performance. If the outcome of the first fine-tuning round does not meet expectations, additional annotated samples can be incrementally added. In such cases, to avoid catastrophic forgetting, all available annotated data should be combined and the model re-fine-tuned starting from the original pre-trained weights, rather than continuing from the previously fine-tuned checkpoint.

### 4.3 Base models and fine-tuning procedures

#### 4.3.1 Cellpose (cyto, cyto3, nuclei, etc.)

Cellpose is a general cell segmentation model based on a U-Net architecture with residual connections. It achieves instance segmentation by predicting vector flows originating from cell centers. It has officially released multiple pre-trained models (such as cyto, cyto3, and nuclei) for different tissue types and staining methods.

CSRefiner integrates an automated fine-tuning process based on Cellpose’s official training interface ‘cellpose.train’. Users can quickly load training data by providing the path file of the image and the corresponding mask pairs. CSRefiner supports grouping based on the sample identifier (SN) in the image file name and uses an optional ratio to split the data into training and validation sets to ensure balanced coverage of each sample during training.

Training utilizes GPU acceleration by default. The optimizer is stochastic gradient descent (SGD) with an initial learning rate of 0.1 and weight decay of 1e-4. The number of training epochs can be configured based on task requirements. After each epoch, the training and validation losses are recorded and saved as JSON log files. Additionally, the loss curve is automatically generated to help users evaluate training progress and model convergence.

The model fine-tuning script is integrated into the unified command-line interface (CLI), supporting one-click switching of different Cellpose pre-trained models. Upon completion, the model weights and log files are automatically saved to a timestamped directory, facilitating version control and reproducibility of experimental results.

#### 4.3.2 Cellpose-SAM (cpsam)

Cellpose-SAM is a new architecture introduced in Cellpose 4.0, which combines the image encoder capabilities of the Segment Anything Model (SAM) released by Meta AI and the flow field prediction mechanism of Cellpose for segmenting cell images. Unlike the original SAM, which relies on a prompt-based decoder, Cellpose-SAM discards the decoder entirely and retains only the pure visual encoder. This encoder directly outputs the vector flows required by Cellpose, making it more suitable for instance segmentation tasks in dense biological images.

In CSRefiner, we provide an independent training script adapted to Cellpose-SAM. The training process is consistent with the previous version of Cellpose, including key steps such as data loading, SN grouping, and loss recording. Users can complete model customization without in-depth understanding of the underlying architecture. Notably, following official recommendations, the Cellpose-SAM training script uses a smaller batch size (1) and a lower learning rate (1e-5) to ensure training stability and robustness.

This architecture has significant advantages in areas with blurred edges and densely populated cells, and is particularly suitable for segmentation fine-tuning scenarios of highly heterogeneous tissue images such as spatial transcriptomes.

#### 4.3.3 StarDist

StarDist is a nucleus segmentation framework tailored to microscopy images, modeling objects as star-convex polygons rather than binary masks. Its backbone is a U-Net encoder–decoder that extracts multi-scale features, followed by two parallel output heads: one predicts object probabilities, and the other predicts radial distances along predefined angles to reconstruct polygons. Final instances are selected through non-maximum suppression, ensuring one unique polygon per nucleus.

CSRefiner incorporates a customized fine-tuning script for StarDist. Notably, the official StarDist framework lacks explicit recommendations for selecting training parameters and does not support the integration of an early stopping function into its training pipeline—two critical limitations that hinder the determination of optimal training epochs. To address this challenge, a single-epoch iterative training strategy was developed. In detail, the training process is executed in iterations of one epoch each, with the loss value recorded at the conclusion of every individual epoch. This design enables real-time dynamic monitoring of the validation set loss; once the model reaches a converged state (characterized by stable or non-decreasing validation loss, patience=30), the training process can be terminated prematurely, thereby simulating the functionality of a standard early stopping mechanism. Users can initialize models from scratch, from official pre-trained weights, or from previously saved checkpoints.

For fluorescent nuclear stains (e.g., DAPI, ssDNA), images are preprocessed as single-channel grayscale with percentile-based intensity normalization; for H&E images, three-channel RGB input is supported. As with other models, logs and weight files are version-controlled, and training curves are automatically generated to assist performance evaluation.

#### 4.3.4 U-Net

CSRefiner integrates a U-Net-based segmentation model designed specifically for biomedical images. It is based on the BCDU-Net D3 architecture and introduces ConvLSTM units in the decoder path to enhance spatial context modeling. With a three-level densely connected decoding structure, U-Net significantly improves segmentation performance on irregular or blurry cell boundaries, making it particularly suitable for tissue images with complex staining patterns or high background noise.

The U-Net fine-tuning process in CSRefiner is implemented based on TensorFlow 2.x. The training process uses a custom DataGenerator class to load and augment data, and the standard fit() interface provided by Keras is used to complete the training process. Preprocessing steps differ based on the staining type. For H&E images, color normalization and intensity normalization are applied using Contrast Limited Adaptive Histogram Equalization (CLAHE). For ssDNA&DAPI images, grayscale conversion, percentile-based intensity normalization, and adaptive histogram equalization are performed.

The model training uses the Adam optimizer and binary cross-entropy loss function. The ReduceLROnPlateau strategy is used to dynamically adjust the learning rate, and early stopping (patience=30 epochs) is implemented to prevent overfitting. The training and validation losses are recorded after each round of training, and a learning curve is plotted to assist analysis.

Users can either train the model from scratch or fine-tune it using a pre-trained .hdf5 weight file. All training outputs (model weights, log files) will be saved uniformly to support reproducibility and downstream inference.

After obtaining a fine-tuned segmentation model, users can apply it to perform inference on complete tissue images or a batch of unannotated image patches to generate high-quality cell segmentation results. CSRefiner provides a unified and efficient inference interface that supports large-scale image processing while ensuring compatibility with downstream spatial transcriptomics analysis pipelines.

### 4.4 cgef file generation

cgef is a widely adopted file format in spatial transcriptomics, designed to integrate gene expression matrices with cell segmentation results. CSRefiner incorporates a dedicated utility for generating ‘.cgef’ files by combining raw expression matrices (‘.gem.gz’ or ‘.gef’ formats) with segmentation masks, leveraging the ‘cgef_writer_cy.generate_cgef’ function from the gefpy library for the core file generation process. The principle is to map each expression point to its corresponding cell segmentation region and associate the segmented cell mask with the spatially resolved expression matrix. For each cell instance, the gene counts from all expression points within its mask are aggregated to generate a cell gene expression matrix annotated with the cell centroid coordinates.

The workflow comprises three key steps. First, input expression matrices undergo standardization: ‘.gem.gz’ files are converted to ‘.gef’ format using an integrated wrapper module. Second, segmentation masks are validated against matrix dimensions, with automatic resizing applied when necessary to ensure alignment with the expression grid. Third, the system classifies masks as either semantic or instance-based, performing conversion of instance masks to semantic format when required before invoking the final ‘.cgef’ generation process.

Output files are generated in blocks (default size: 256 × 256 pixels) to accommodate large-scale datasets while minimizing memory consumption. The utility automatically cleans up intermediate files and maintains comprehensive logs for full process traceability. This streamlined pipeline enables reproducible integration of cell segmentation data with transcriptomic quantification, facilitating downstream analytical tasks including cell annotation, clustering, and differential expression analysis.

### 4.5 Cell segmentation performance evaluation metrics

In order to comprehensively evaluate the performance of different models and fine-tuning schemes in the cell instance segmentation task, we use five classic evaluation indicators: Precision, Recall, F1 Score, Jaccard Index and Dice Coefficient. These indicators focus on measuring the overlap and matching quality between the predicted mask and the true mask, and are suitable for quantitative comparison of the generalization ability of the model in small sample training scenarios.

Precision: measures how many of the positive samples predicted by the model are real cell instances. A high precision indicates that the model prediction is more conservative and has fewer false positives (FP).

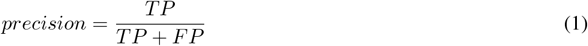

Recall: indicates how many of the real cell instances are correctly identified by the model. A high recall indicates that the model is more capable of finding all targets and has fewer missed detections (FN).

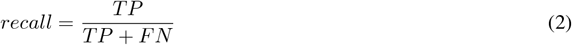

F1 Score: the harmonic mean of precision and recall, which comprehensively reflects the model’s precision and recall capabilities and is an important measure of the model’s overall performance.

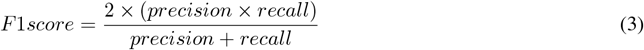

Jaccard Index: measures the ratio of the intersection and union between the predicted mask and the true mask. It is a commonly used overlap indicator in instance segmentation tasks. The higher the value, the more accurate the prediction.

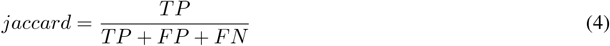

Dice Coefficient: similar to Jaccard, it also measures the degree of overlap between masks, but is more sensitive to small targets and is often used to evaluate fine structure segmentation in the field of biological images.

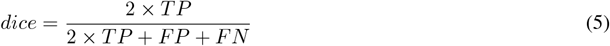

Among them, TP (True Positive) indicates the number of cell instances correctly identified by the model, FP (False Positive) indicates false positives, and FN (False Negative) indicates missed positives. We use the IoU matching strategy to match the predicted mask with the true mask one by one, and set a threshold (default is IoU > 0.5) to determine whether the match is established.

All metric results are computed based on the model’s predicted masks and the corresponding groundtruth masks, and are recorded in .xlsx files for tracking and reproducibility. We use Python’s matplotlib[33] library to visualize these metrics, including bar charts and box plots, which clearly illustrate the differences in average performance and inter-sample stability across models.

### 4.6 Quantification of cellular morphological and spatial metrics

All cellular images were acquired in TIFF format, accompanied by corresponding binary segmentation masks that delineate the boundaries of individual cells. Masks were converted to boolean arrays (where True values represented cellular regions and False values represented the background) to enable precise region-of-interest (ROI) analysis. Image preprocessing and metric calculation were performed using Python[34] with core libraries including scikit-image[35] for image processing, NumPy[36] for numerical computations, pandas[37] for data organization, and SciPy[38] for spatial distance calculations.

A custom CellMetricsCalculator class was developed to automate the extraction of six key cellular metrics, encompassing spatial arrangement, density, morphological complexity, and boundary integrity. For each image-mask pair, cellular regions were first labeled using connected-component analysis, and region properties were extracted to quantify area, perimeter, and centroid coordinates of individual cells. Detailed calculations for each metric are described below.

#### 4.6.1 Mean Distance

Specifically, the average nearest-neighbor distance quantifies the mean spatial separation between each cell and its closest neighboring cell, reflecting the overall dispersion of cells within the field of view.

For all labeled cells, centroid coordinates (*x*_*i*_, *y*_*i*_) (where *i* = 1, 2, …, *N* and *N* is the total number of cells) were computed. A pairwise Euclidean distance matrix between all centroids was generated using the formula:

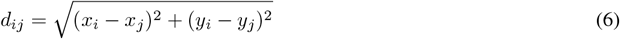

Where *d*_*ij*_ is the Euclidean distance between the centroids of cell *i* and cell *j*. Diagonal elements of the distance matrix (representing self-distance) were set to infinity, and the minimum distance for each cell (*d*_*min,i*_ = min(*d*_*i*1_, *d*_*i*2_, …, *d*_*iN*_, *i* = *j*)) was identified. The average nearest-neighbor distance was calculated as the arithmetic mean of all *d*_min,*i*_:

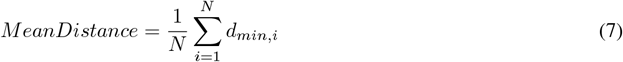

#### 4.6.2 Compactness

Compactness characterizes the degree of cell packing by normalizing nearest-neighbor distance to cellular size, with higher values indicating looser packing and lower values indicating tighter packing.

Average cellular area *Ā* was computed as the mean area of all labeled cells:

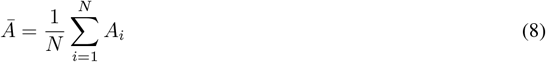

Where *A*_*i*_ is the area (in pixels^2^) of cell *i*. Compactness was derived by normalizing the squared average nearest-neighbor distance (to account for area units) by the average cellular area:

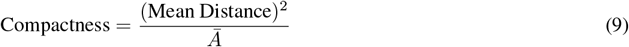

#### 4.6.3 Density

Cell density quantifies the number of cells per unit area, reflecting the overall confluency and population density of the cellular monolayer.

Total cellular area (*A*_*total*_) was calculated as the sum of pixels in the binary mask (i.e., the total area occupied by all cells). Cell density was computed as the ratio of the total number of cells (*N*) to *A*_*total*_:

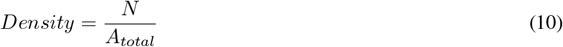

#### 4.6.4 Distribution

Distribution assesses the heterogeneity of local cell density, with higher values indicating uneven spatial distribution (e.g., formation of cell clusters) and lower values indicating uniform dispersion.

For each cell *i*, Euclidean distances to all other cells were calculated, sorted, and the distances to the *k* = 3 nearest neighbors (excluding the cell itself) were summed 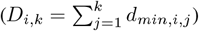. Local density for cell *i* was defined as 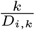 (inverse of the average distance to *k* neighbors). Population-level mean (*µ*_*local*_) and standard deviation (*σ*_*local*_) of local densities were computed. Clustering was derived as the coefficient of variation (CV) of local densities:

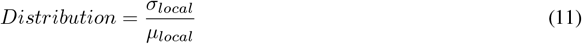

#### 4.6.5 Shape Complexity

Shape Complexity describes the irregularity of cellular shapes, with higher values indicating more complex (e.g., branched, irregular) cell boundaries and lower values indicating simpler (e.g., circular, oval) shapes. This metric is derived from the relationship between cellular perimeter and area, adapted for population-level analysis.

For each cell *i*, perimeter (*P*_*i*_) and area (*A*_*i*_) were extracted from region properties. Population-level averages of squared perimeter 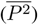 and area 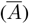 were computed:

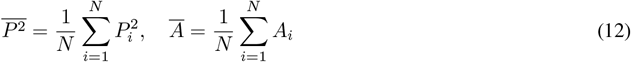

Morphological complexity was defined as the ratio of 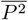 to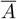:

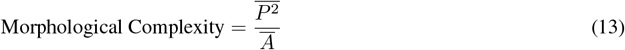

#### 4.6.6 Edge Contrast

Edge Contrast quantifies the contrast between cellular edges and the background, reflecting the sharpness of cell boundaries in the original image.

RGB images (if applicable) were converted to grayscale via intensity averaging across color channels. A gradient magnitude image was generated using the Sobel operator (filters.sobel), which highlights edge regions by measuring intensity changes across the image. Gradient values were partitioned into two regions: Cell-edge gradient: Gradient values within the binary mask (i.e., edges of cellular regions), with mean intensity *µ*_*cell*_. Background gradient: Gradient values outside the binary mask (i.e., background edges), with mean intensity *µ*_*bg*_. Boundary clarity was computed as the ratio of *µ*_*cell*_ to *µ*_*bg*_:

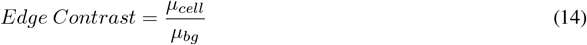

### 4.7 Downstream transcriptomic analysis

To assess the biological impact of improved segmentation, downstream transcriptomic analyses were performed using the Stereo Data Analysis Solution (SDAS, available at https://github.com/STOmics/SDAS). SDAS is a dedicated platform for advanced spatial transcriptomics analysis, providing modules for cell annotation, differential expression testing, and functional enrichment. The resulting cgef file output by the CSRefiner pipeline was converted to an AnnData object using Stereopy[39, 40] before being input into SDAS.

#### 4.7.1 Cell type annotation

Cell type annotation was performed with the cell2location[23] module within SDAS. For each segmented cell, annotation scores across all reference types were computed, and the cell was assigned to the type with the highest posterior probability. Annotation confidence was quantified by the distribution of assignment scores, and population-level shifts in score distributions between pre-trained and fine-tuned results were compared. Annotation outputs were visualized in spatial maps and used as the basis for downstream analyses.

#### 4.7.2 Differential expression gene analysis

Differential expression gene (DEG) testing was conducted for each annotated cell type using the Wilcoxon rank-sum test implemented in SDAS. Genes with false discovery rate (FDR) *<* 0.05 and absolute log-fold-change *>* 1 were defined as significant DEGs. To capture global trends, we quantified the number of significant DEGs per cell type before and after fine-tuning, with particular focus on the six most enriched cell populations identified in the hippocampal region (e.g., Ext_Hpc_DG1, CA1–CA3 subfields). Comprehensive DEG tables, including both raw statistical results and filtered sets, were generated for reproducibility.

#### 4.7.3 Functional enrichment analysis

Significant DEGs were further analyzed for functional relevance using GO Biological Process (2023) annotations via Enrichr[41]. Upregulated and downregulated gene sets were tested separately, and enrichment results were exported as tabular files and visualized through multiple modalities, including bubble plots (summarizing gene counts, rich factors, and adjusted significance), waterfall plots (depicting representative genes within top enriched pathways), and, where appropriate, heatmaps of gene expression patterns across GO terms. These analyses highlighted biological pathways more comprehensively recovered after fine-tuning, thereby linking improved segmentation to enhanced interpretability of spatial transcriptomic data.

## 5 Data availability

The two datasets that support the findings of this study have been deposited into CNGB Sequence Archive (CNSA) of the China National GeneBank DataBase (CNGBdb) with accession number CNP0007731: https://db.cngb.org/search/project/CNP0007731/. Additionally, we have uploaded these datasets to Zenodo[42].

## 6 Code availability

Project home page: https://github.com/STOmics/CSRefiner

## 7 Authors’ Contributions

Project administration and supervision: Mei Li, Ying Zhang, Sha Liao, Ao Chen

Algorithm development and implementation: Can Shi

Data provision: Yumei Li, Jing Guo, Qiuling Chen, Tingting Cao

Data selection and processing: Can Shi, Yumei Li, Ying Zhang

Data annotation: Can Shi

Code Testing: Can Shi

Manuscript writing and figure generation: Can Shi, Ying Zhang

Manuscript review: Can Shi, Mei Li, Ying Zhang

## 9 Acknowledgments

We thank the China National GeneBank for providing data storage support for this study and the National Key R&D Program of China (2022YFC3400400) for funding. We also thank Zhonghan Deng, Manqi Liang, Jiaxue Chen and Zhihan Ren for their assistance.

