## Supplementary Fig. 1 for "CSRefiner: A lightweight framework for fine-tuning cell segmentation models with small datasets"

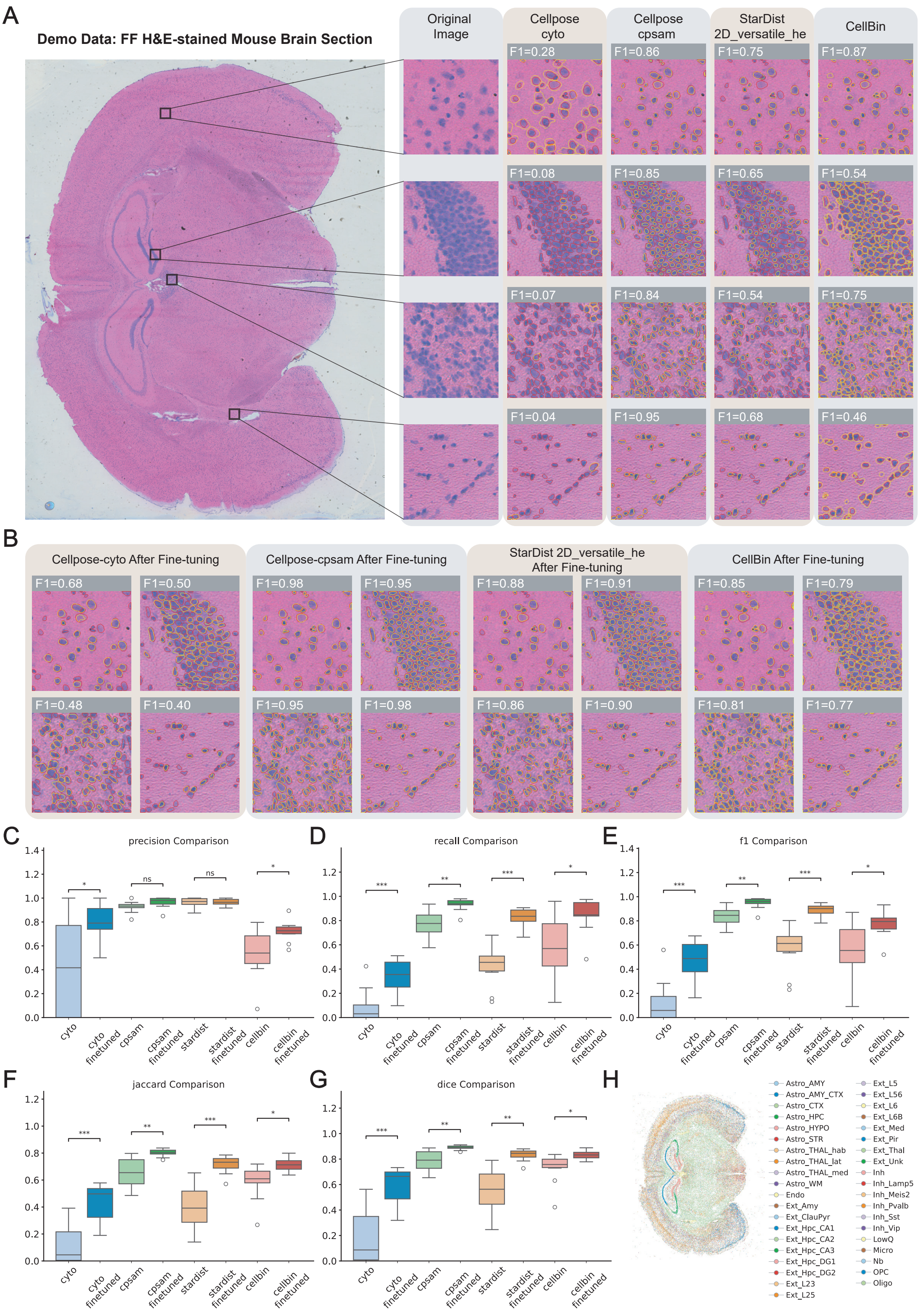


Figure S1: **Performance of pre-trained and fine-tuned segmentation models on FF H&E-stained mouse brain section.** (A) Representative segmentation results from four pre-trained models (Cellpose-cyto, Cellpose-cpsam, StarDist-2D_versatile_he, and CellBin) for FF mouse brain tissue with H&E-stained. Red contours indicate ground-truth manual annotations, and yellow contours indicate model-predicted segmentation boundaries; the corresponding F1 scores are labeled for quantification. (B) Segmentation results after fine-tuning with CSRefiner, corresponding to the regions shown in panel A. (C–G) Quantitative evaluation of segmentation performance before and after fine-tuning across four representative models (Cellpose-cyto, Cellpose-cpsam, StarDist, and CellBin). Boxplots show improvements in (C) precision, (D) recall, (E) F1 score, (F) Jaccard index, and (G) Dice coefficient. Significance indicators (ns, *, **, ***) denote P values $> 0.05$, $< 0.05$, $< 0.01$, and $< 0.001$, respectively. (H) Spatial maps of cell type annotations generated by cell2location using cgef matrices from the fine-tuned cpsam model.
